# Put the frog in water: simple methods for improving individual identification as demonstrated with the agile frog

**DOI:** 10.1101/2025.02.25.640084

**Authors:** Edina Nemesházi, Zsanett Mikó, Nikolett Ujhegyi, Andrea Kásler, Nadine Lehofer, Veronika Bókony

## Abstract

Identification of individuals across time and space is required for investigating numerous evolutionary-ecology and conservation-related questions, and photo-based identification is commonly used for a broad taxonomic range. Systematic comparisons of available photo-matching software have been published for various taxa, and success of such software may greatly depend on image quality as well as the focal body parts. Yet, suitability assessments of images captured with different methods are missing. Here we tested the hypotheses that individual identification by colour patterns can be facilitated by taking into account the natural medium surrounding the animals and the natural body posture they tend to take. We developed new photography methods to enable individual identification by whole-body assessment using the agile frog *(Rana dalmatina)*, a species with common yet so-far neglected melanin-based pattern on its limbs and back. We compared the reliability of different photography methods for computer-assisted identification in the HotSpotter software as well as for observers operating it. We found that photographing either hand-restrained frogs with towel-dried skin, or frogs moving freely in clean water enabled comparison of the dorsal surface of the whole body including the hind legs, and HotSpotter identified matching images at rates similar to the relatively more successful of previously published anuran studies (>92% listed with rank≤10). By contrast, submerging hand-restrained frogs in water significantly improved identification: images of re-captured individuals were always ranked as the most likely match. We attribute this outstanding performance to the combination of advantageous effects of in-water light refractions that improve the visibility of melanin patterns, and uniform body postures facilitating comparison across individuals. Observers in general successfully identified matching images and ruled out non-matching ones, but some mistakes were recorded when images featured freely moving frogs. The photography methods developed in this study should be easily adapted to most frog and toad species for reliable individual identification. Our study highlights that taking into account features of the natural environment of the studied species can improve individual identification by photographs. Because such methods are non-invasive and inexpensive, they should be especially beneficial for population monitoring programs of endangered species.

**Data availability:** Data are available from the FigShare Repository [DOI will be assigned upon acceptance for publication]. Further supporting information for methods and results is provided as Supplement to this manuscript.

## Introduction

Capture-recapture methods enable the assessment of survival rate, growth rate, life span and further questions of evolutionary-ecology studies and conservation management (Canessa et al., 2019; Sarasola-Puente et al., 2011). Individual identification (hereafter ‘ID’) is therefore key in population monitoring, and it can be performed either using artificial tagging methods (such as injecting transponders or colourful elastomers under the skin; Donnelly et al., 1994), genotyping (Lukacs & Burnham, 2005), or exploiting natural markings (Bohnett et al., 2023; Dunbar et al., 2021; Matthé et al., 2017). In the latter case, individual markings are captured by photography, and computer-assisted image comparison methods can facilitate ID in large populations across multiple years. While artificial tags can be lost (Bendik et al., 2013; Donnelly et al., 1994), macroscopic colour patterns such as spots and stripes (hereafter ‘melanin patterns’) remain largely stable at least during adult life (Price et al., 2008; Rojas et al., 2023), providing suitable resources for non-invasive or minimally invasive ID. Photo-based ID has been used in various taxa including mammals, reptiles and amphibians, and systematic comparisons of test databases have been conducted to assess the performance of different software that aim to reduce the time requirement of photo comparison (Bohnett et al., 2023; Burgstaller et al., 2021; Dawson et al., 2021; Matthé et al., 2017; Morrison et al., 2016). While much attention has been paid to software performance, capture-recapture studies could benefit from optimized photography methods. Because species-specific natural markings evolved to be visible or provide camouflage in specific habitats combined with natural behaviour (Gomez & Théry, 2004; Rojas, 2017), body posture and the environment including the surrounding medium may both influence pattern recognition.

The effect of different surrounding media on pattern visibility may be especially relevant for amphibians, a taxonomic group highly threatened by various global changes (Luedtke et al., 2023), because individuals of most species use terrestrial as well as aquatic habitats, changing with life stages and reproductive vs. non-reproductive periods. Because adult amphibians are typically solitary throughout the year but aggregate in aquatic habitats for breeding, melanin patterns playing a role in recognizing territory neighbours, choosing mates and other types of visual communication might have evolved to be most visible in water in various species. Previous studies used photographs of anurans out of water either restrained by hand (Kim et al., 2017; Lama et al., 2011), placed on their back to enable the assessment of ventral patterns (Aevarsson et al., 2022; Caorsi et al., 2012), or sitting in a natural posture on a dry surface (Burgstaller et al., 2021; Davis et al., 2020; Morrison et al., 2016; Patel & Das, 2020). While this latter approach might be less stressful to the animals, it is not suitable for assessing patterns across the legs, because those are usually tightly folded on both sides of the body. In contrast, anurans tend to take a body posture with more stretched hind legs in water environment (Fig. 1); and exploiting this natural behaviour might improve ID without having to restrain the animals. Furthermore, differential light refractions in water compared to air might improve the visibility of colour patterns. Image quality issues regarding light conditions and reflections on frog or toad skin have been mentioned in various studies either as nuisances to avoid or as a suspected reason for reduced matching success in photo-based ID (Dawson et al., 2021; Kim et al., 2017; Matthé et al., 2017; Morrison et al., 2016; Patel & Das, 2020), but to our knowledge no study attempted to optimise and systematically compare the suitability of photography methods in this taxon so far.

**Fig. 1.**
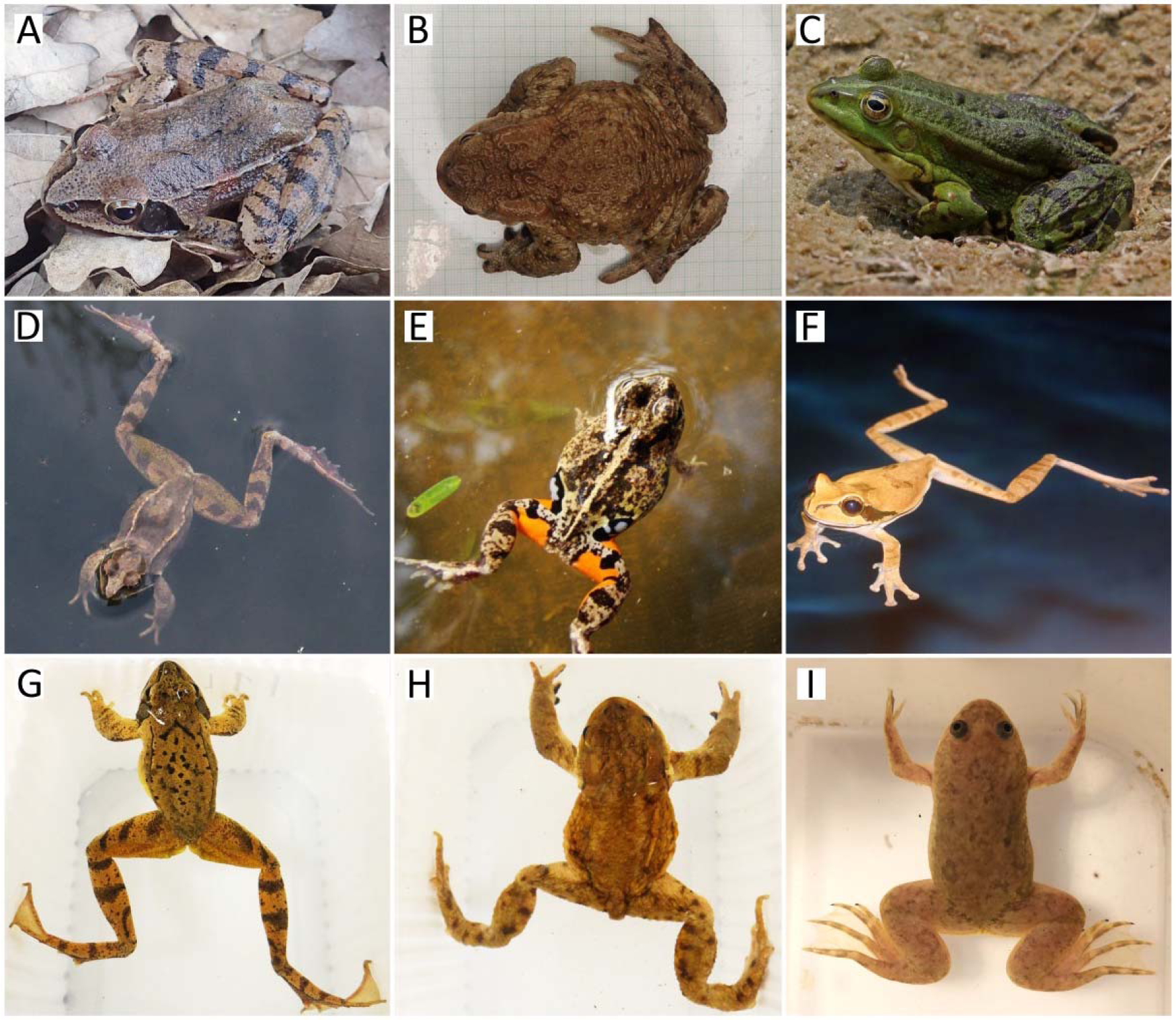
Examples of body postures of frogs and toads on land (A-C) and in water (in natural environment: D-F, or in an unfamiliar white box: G-I). While legs are typically tightly folded on land, body postures in water often provide better visibility of them. Species: agile frog (*Rana dalmatina*; A, D, G), common toad (*Bufo bufo*; B, H), marsh frog (*Pelophylax ridibundus*; C), Colombian four-eyed frog (*Pleurodema brachyops*; E), masked tree frog (*Smilisca phaeota*; F), and African clawed frog (*Xenopus laevis*; I). Photo credit: Nikolett Ujhegyi (A, B, H), Charles J. Sharp (C, F), Edina Nemesházi (D, G), Luis Alberto Rueda (E) and Zsanett Mikó (I).

Most pattern-recognition software has strict requirements for image layout, therefore may only be used on a well-defined area of the body and often require complex image pre-processing (Burgstaller et al., 2021; Matthé et al., 2017). Therefore, amphibian photo-based ID studies so far usually focused on a single body part that was deemed to feature suitably variable patterns between individuals of the focal species. Usage of dorsal patterns is more common (Burgstaller et al., 2021; Davis et al., 2020; Dawson et al., 2021; Morrison et al., 2016; Patel & Das, 2020), but ventral patterns (Aevarsson et al., 2022; Caorsi et al., 2012; Matthé et al., 2017), lateral patterns on one hind leg (Lama et al., 2011) and lateral lines (Kim et al., 2017) have also been used with varying success (Table S1). However, in anurans the legs often feature at least as, if not more prominent melanin patterns than the back (e.g. majority of species in Ranidae, the large family of ‘true frogs’; but see further examples on Fig. 1). Developing a photo-based identification protocol that enables the exploitation of melanin patterns across both the back and dorsal surface of the legs could therefore be most beneficial; and increasing the amount of collected data may facilitate differentiation between similar-looking individuals.

In the present study, we tested different photography methods for melanin-pattern comparison across the dorsal surface of the body and legs of agile frogs (*Rana dalmatina*). Our aim was to assess the suitability of different image types for computer-assisted ID while enabling observers to compare as much of the body-wise patterns as possible. We chose the freely accessible HotSpotter software (Crall et al., 2013), because it proved to be suitable for whole-body comparison in a mammal (Bohnett et al., 2023), and in an anuran species as well (using images with limited hind-limb visibility; Burgstaller et al., 2021). Over the course of two study years, we created a total of 12 databases (six for males and six for females) consisting of images featuring frogs either positioning themselves freely in water or restrained by hand and photographed either with dried skin or being submerged in water. We assessed if the animal’s pose (i.e. restrained or free), surrounding medium (air or water), sex, and database size affected ID success across hundreds of individuals.

## Methods

### Animal collection

We captured free-ranging agile frogs during the breeding season in 2023 and 2024 at a breeding pond near Budapest (47°33’04.3•N, 18°55’36.0"E), along a c.a. 1.5-m tall drift fence surrounding the pond, that was dug 20-cm deep underground with pitfall traps both inside and outside. The fence was standing between 03 March and 05 April in 2023, and between 06 February and 18 March in 2024. We checked the traps every morning for frogs arriving to breed or leaving the pond (hereafter, we will refer to collecting a frog from a trap as a ‘capture event’). Only a subset of arriving frogs was captured in 2023 because a number of males arrived before fence building, and strong wind temporarily destroyed the fence on 11 March. In both years, only a subset of the arriving individuals was re-captured upon leaving, because some frogs managed to escape the fence using climbing surfaces available in the inside, and some may have stayed in the pond longer. The animals were carried to our nearby (ca. 700 m) experimental station in Julianna-major at the Plant Protection Institute, Budapest for photographing, and each individual was subsequently placed to that side of the fence where it was heading at capture. We assigned a unique identification number to each capture event that was photographed together with each frog.

### Photography

We photographed agile frogs in teams of two people, where one person captured photographs (hereafter ‘photographer’) and the other handled the frogs (hereafter ‘animal handler’). Across the study, we used a Sony Cyber-shot DSC-HX200V camera (saving jpg files in 350 dpi resolution), except for 5 ‘dry-restrained’ images of arriving females in 2023, that were captured with an Olympus Tough TG-4 camera). We captured three image types (i.e. using three photography methods) in total. The ‘dry-restrained’ image type (upper panel on Fig. 2) featured hand-restrained frogs photographed after their skin was gently dried with towel or paper towel, and their hind legs being gently pulled back by the ankles (aiming for a relatively uniform body posture). When capturing the ‘water-free’ image type (Fig. 3 A), frogs were placed into water where they chose their body posture freely (but posture changes were gently initiated by hand if needed, for example if the legs were tightly folded or the animal stood vertically). These two image types were captured in both 2023 and 2024. Additionally, based on our results and experiences obtained in 2023, in 2024 we decided to capture a third image type as well (i.e. ‘water-restrained’, Fig. 3 B), where frogs were hand-restrained in water (see Fig. 2 for illustration). Learning from technical issues noted in 2023 (Fig. S1), we instructed photographers in 2024 to aim to keep the camera’s lens parallel to the back of each frog and avoid direct light exposure (i.e. lean above the animal or use an umbrella fixed on the table to obscure the ceiling light, and never use the camera’s flash; Fig. 3 C). We also instructed animal handlers to avoid obscuring the lower-leg patterns by their fingers, avoid twisting or stretching the hind legs to a straight posture, clean each animal in a water bath before placing it into the water where frogs are photographed, and change the latter water as needed to minimise floating debris and bubbles. Building on our experiences, we provide a detailed description of the best photography practices in Supplement Section II.

**Fig. 2.**
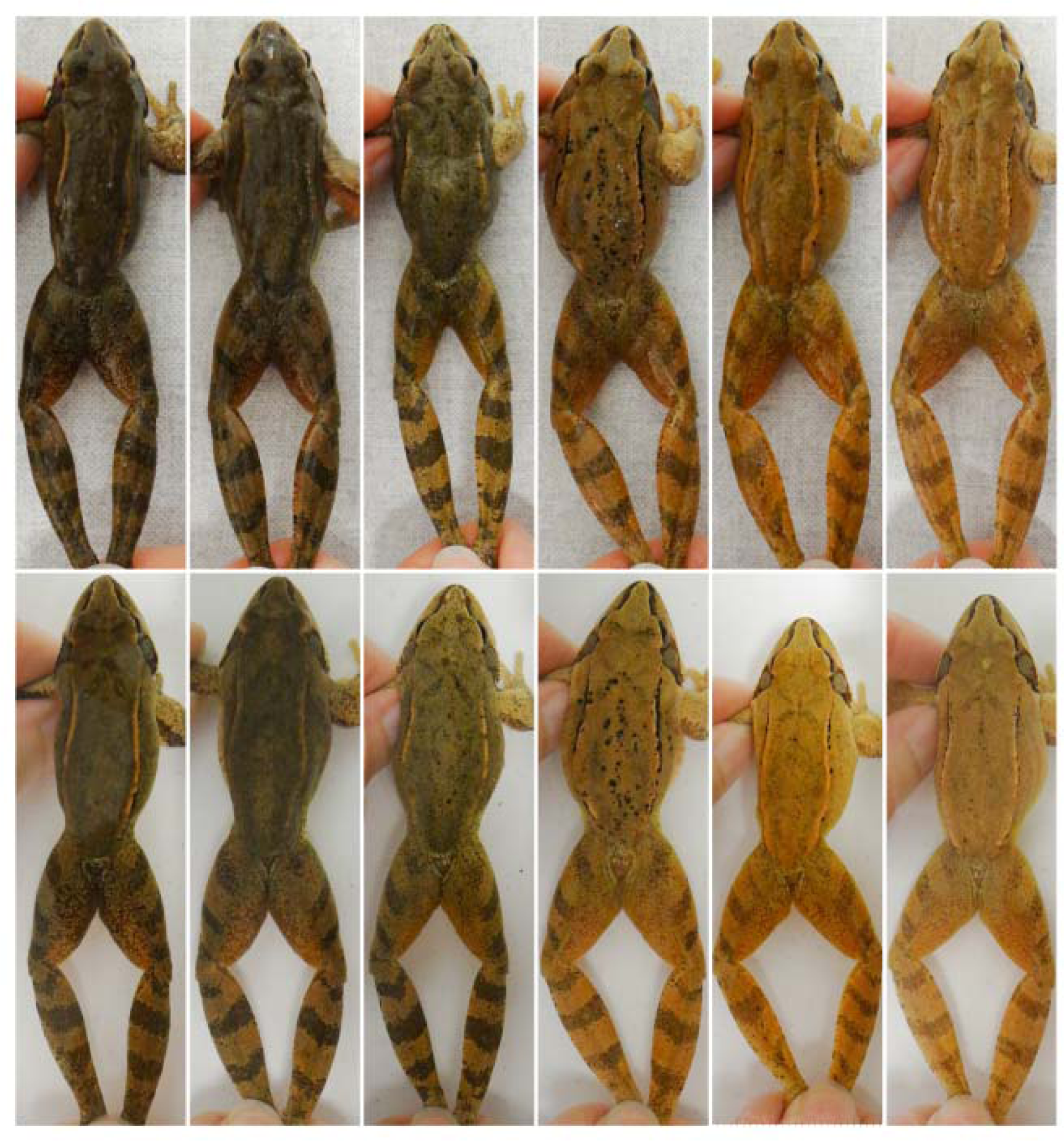
Agile frog individuals with various skin tones photographed after paper-towel drying (top) and under water (bottom). For each individual, the two images were captured a few seconds apart, using the same light source, while avoiding direct-light exposure to minimise reflections. Note how submerging in water decreases skin glossiness and shadows caused by uneven body surface, overall improving the perception of contrast between skin tone and melanin patterns. For illustration purposes on this figure, we applied the same exposure curve across all images in RawTherapee (referred to as ‘baseline’ on Fig. S2), and cropped them subsequently.

**Fig. 3.**
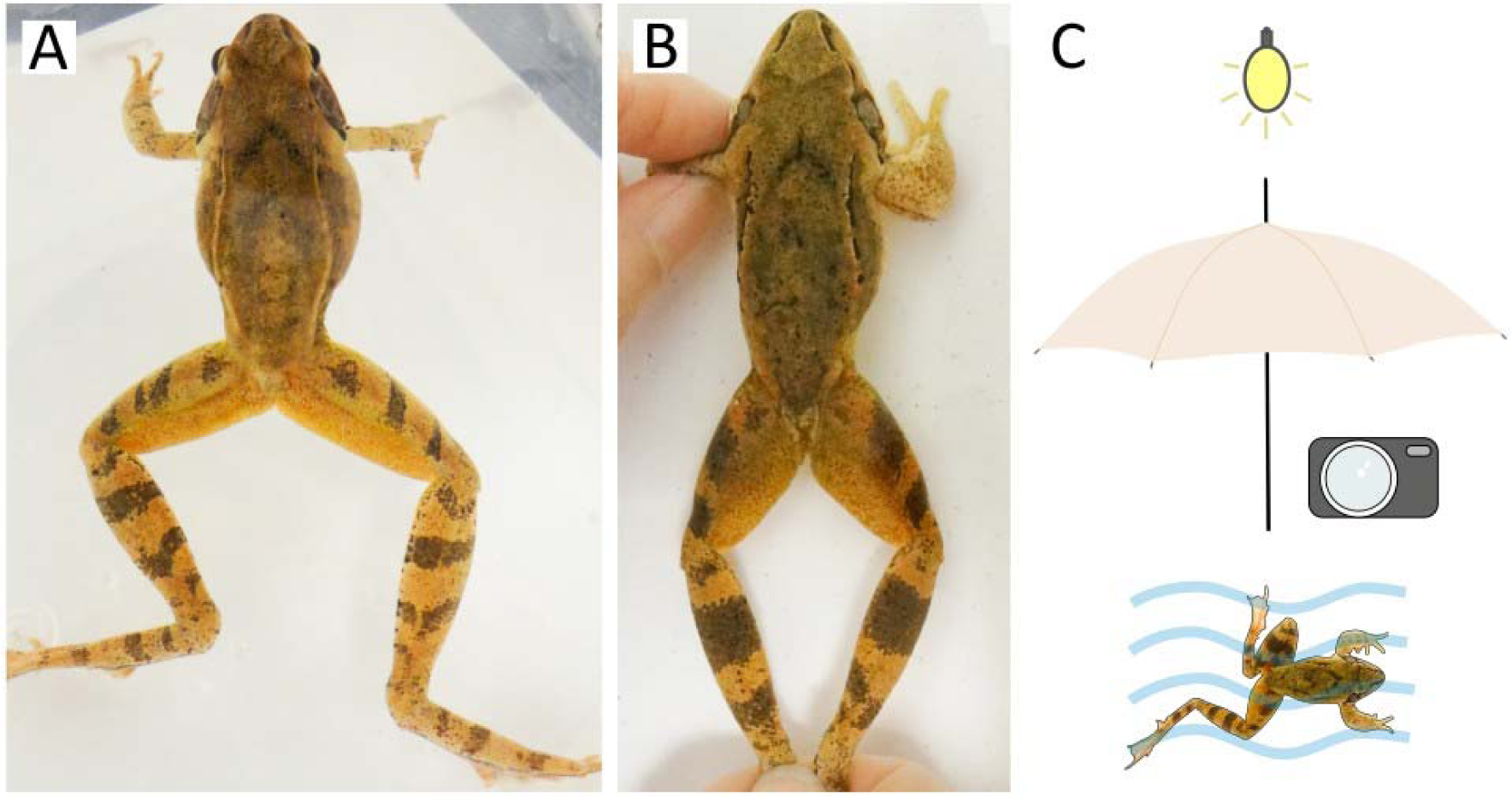
Fine-tuned photography of frogs submerged in water in 2024. The two poses are shown in water medium: ‘free’ (A) and ‘restrained’ (B). The images were cropped to indicate the approximate coverage of the frog body that was used for ID by HotSpotter. Panel C shows visual representation of key methodological components for both image types: frogs were photographed in water, while direct light exposure was physically prevented.

### Image processing

We manually screened all images and chose the ones deemed most suitable for identification from each capture event. Suitability was judged based on the following criteria: ideally, the animal’s back and hind legs should be in focus, reflections should not obscure any potential melanin patterns, and the largest possible surface of both the back and hind legs should be visible on the image. Similarly to other anurans, the hue and darkness of the skin is diverse and can change substantially in agile frogs, even within the period of a few weeks between arriving to and leaving the pond. Therefore, to increase the contrast between skin tone and melanin patterns, we applied exposure curves in RawTherapee 5.9 (The RawTherapee Team, 2022). While these were set image by image in 2023, we used one of five pre-saved exposure curves in 2024 that we optimized for different skin tones under the here-applied light conditions (Fig. S2). In a few cases we manually altered these curves to further improve melanin-pattern visibility based on personal judgement in 2024 as well. We imported images into HotSpotter (Crall et al., 2013) databases, where we set ‘chips’ (i.e. the region of interest on each image in HotSpotter) featuring the entire body of the frog (but toes may be excluded to gain a better zoom); and set the orientation of the animals so the nose pointed to the left, and the vertebral column was horizontal. Because our aim was to gain as much information from each image as possible, we did not exclude the handlers’ fingertips from the ‘chips’ at the cost of cropping out visible parts of the frog body.

First we created 10 HotSpotter (Crall et al., 2013) databases separated by image type (‘water-free’, ‘water-restrained’ or ‘dry-restrained’), year (note that ‘water-restrained’ images were only captured in 2024) and sex (because females undergo substantial body-shape changes during egg laying and might also feature behavioural differences, potentially affecting melanin-pattern visibility). When designing the databases, our aim was to enable comparison of the performance of HotSpotter as well as observers between image types. For this purpose, within each year for each sex, the above-mentioned HotSpotter databases consisting of different image types were identical in size and featured one image per capture event across the same set of capture events (i.e. same set of female captures from the given year or same set of male captures from the given year across the different image types). The detailed list and respective sizes of databases are shown in Table 1). All databases created in 2023 were of similar size. In 2024, we created larger databases from the in-water photos, but for comparability with the ‘dry-restrained’ database we created one subset database (named ‘water-restrained subset’) featuring ‘water-restrained’ images for each sex with the same sample size as for the ‘dry-restrained’ dataset. Overall, this resulted in a total of 12 HotSpotter databases in our analyses.

**Table 1.**
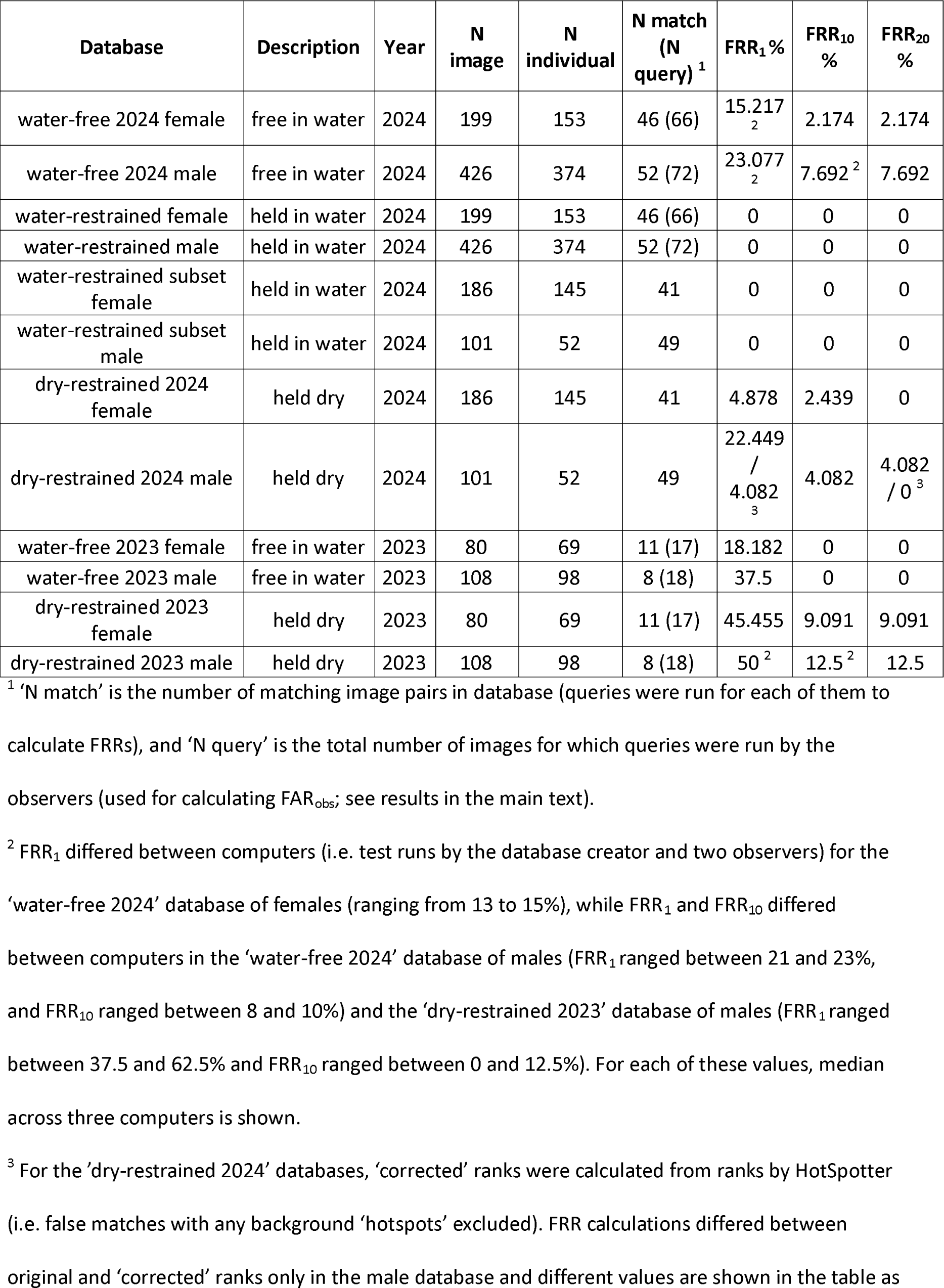
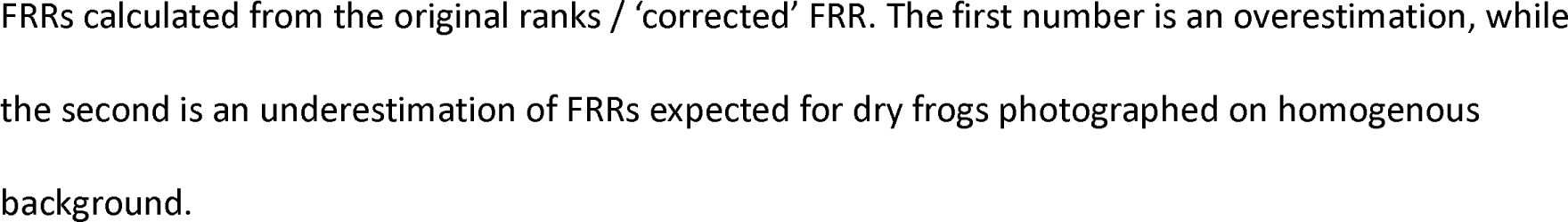
False rejection rates by HotSpotter in different databases.

| Database | Description | Year | N image | N individual | N match (N query) <sup>1</sup> | FRR<sub>1</sub> % | FRR<sub>10</sub> % | FRR<sub>20</sub> % |
| --- | --- | --- | --- | --- | --- | --- | --- | --- |
| water-free 2024 female | free in water | 2024 | 199 | 153 | 46 (66) | 15.217<sub>2</sub> | 2.174 | 2.174 |
| water-free 2024 male | free in water | 2024 | 426 | 374 | 52 (72) | 23.077<sub>2</sub> | 7.692 <sup>2</sup> | 7.692 |
| water-restrained female | held in water | 2024 | 199 | 153 | 46 (66) | 0 | 0 | 0 |
| water-restrained male | held in water | 2024 | 426 | 374 | 52 (72) | 0 | 0 | 0 |
| water-restrained subset female | held in water | 2024 | 186 | 145 | 41 | 0 | 0 | 0 |
| water-restrained subset male | held in water | 2024 | 101 | 52 | 49 | 0 | 0 | 0 |
| dry-restrained 2024 female | held dry | 2024 | 186 | 145 | 41 | 4.878 | 2.439 | 0 |
| dry-restrained 2024 male | held dry | 2024 | 101 | 52 | 49 | 22.449 / 4.082<sub>3</sub> | 4.082 | 4.082 / 0<sup>3</sup> |
| water-free 2023 female | free in water | 2023 | 80 | 69 | 11 (17) | 18.182 | 0 | 0 |
| water-free 2023 male | free in water | 2023 | 108 | 98 | 8 (18) | 37.5 | 0 | 0 |
| dry-restrained 2023 female | held dry | 2023 | 80 | 69 | 11 (17) | 45.455 | 9.091 | 9.091 |
| dry-restrained 2023 male | held dry | 2023 | 108 | 98 | 8 (18) | 50<sup>2</sup> | 12.5<sup>2</sup> | 12.5 |

<sup>1</sup> 'N match' is the number of matching image pairs in database (queries were run for each of them to calculate FRRs), and 'N query' is the total number of images for which queries were run by the observers (used for calculating FAR<sub>obs</sub>; see results in the main text).
<sup>2</sup> FRR<sub>1</sub> differed between computers (i.e. test runs by the database creator and two observers) for the 'water-free 2024' database of females (ranging from 13 to 15%), while FRR<sub>1</sub> and FRR<sub>10</sub> differed between computers in the 'water-free 2024' database of males (FRR<sub>1</sub> ranged between 21 and 23%, and FRR<sub>10</sub> ranged between 8 and 10%) and the 'dry-restrained 2023' database of males (FRR<sub>1</sub> ranged between 37.5 and 62.5% and FRR<sub>10</sub> ranged between 0 and 12.5%). For each of these values, median across three computers is shown.
<sup>3</sup> For the 'dry-restrained 2024' databases, 'corrected' ranks were calculated from ranks by HotSpotter (i.e. false matches with any background 'hotspots' excluded). FRR calculations differed between original and 'corrected' ranks only in the male database and different values are shown in the table as
FRRs calculated from the original ranks / 'corrected' FRR. The first number is an overestimation, while the second is an underestimation of FRRs expected for dry frogs photographed on homogenous background.

For testing purposes, each individual was represented by maximum two images in each database. We ensured this by first identifying all recaptures in 2023 and 2024 for each sex as described in Supplement Section I. To create the 12 HotSpotter databases for our analyses, we selected capture events for which at least two of the three image types were available and photography methods in general met our pre-defined methodological guidelines (see details on image selection in Supplement Section III). For those individuals that were represented by two images in a database, we assigned one ‘chip’ (corresponding to one image) to be queried in HotSpotter, and the other ‘chip’ as its ‘true match’ that should be found. When running a query, HotSpotter calculated pairwise similarity scores between the ‘chips’ of the query image and all other images in the database, and ranked them accordingly with rank 1 meaning the highest similarity. Each database was run on the database creator’s computer (the same computer across all databases), where we recorded ranks and scores assigned to ‘true matches’. Also, we archived visual results of HotSpotter by saving screen shots of true matching pairs of images as presented by HotSpotter, with and without circles that denoted features identified by HotSpotter as informative similarities between two images (hereafter referred to as ‘hotspots’). Additionally, databases representing each of the three image types were handed over to two out of four database operators (hereafter referred to as observers) who had not previously worked with these databases and used HotSpotter in 2023 for the first time. Observers were instructed to run a query for each of a list of ‘chips’ and systematically screen the resulting image pairs between rank=1 (best) and rank=20 until they found a matching image. These ‘chips’ to be queried featured individuals that belonged to one of two categories: 1) were re-captured, having exactly one matching ‘chip’ in the database that should be found, or 2) had no matching ‘chip’ in the database. The observers were unaware of these categories during the tests. To ensure that overall observer performance with each image type was not influenced by recent experience, within each year, two observers worked first with one image type and subsequently with another, while the other two observers followed the opposite order. While observers tested all four databases in 2023, they did not test the ‘dry-restrained 2024’ and ‘water-restrained subset’ databases in 2024, because they proved to be able to reliably work with the dry image type in 2023 already, and ran queries for the ‘water-restrained’ image type in the larger database versions in 2024.

### Statistical analyses

For each database, we calculated false rejection rate (FRR), which is the number of falsely rejected matches divided by the number of matching pairs present in the database for which queries were performed, expressed as a percentage (Bolger et al., 2012; Jain, 2007). We applied three thresholds: we accepted a match by HotSpotter if it was the image that received the highest rank (FRR_1_), or among the 10 highest ranked images (FRR_10_) or the 20 highest ranked images (FRR_20_), respectively.

‘Dry-restrained’ images in 2024 were photographed on a workbench that had light grey ornamentation on white surface (see the upper panel on Fig. 2). This may lead to overestimation of FRR if false matches received higher similarity scores due to background similarities. To account for this potential effect, we manually checked for each query if false ‘hotspots’ were indicated on the background on those non-matching images that were ranked higher than the matching image. Based on this information, we calculated ‘corrected’ ranks for the true matches by excluding all those higher-ranking false matches where even a single false ‘hotspot’ was indicated on the background, and we used these ‘corrected ranks’ for calculating ‘corrected’ FRR_1_, FRR_10_ and FRR_20_ (i.e. underestimated values of FRR).

To test whether FRR varied systematically by the photographing medium, body pose, sex, year, and database size, we used the data from all databases in a single generalized linear mixed-effects model with binomial error and logit link. To handle repeated measures due to the inclusion of the same individuals into different databases, we treated individual identity as a random factor. The four fixed factors and the fixed covariate (database size standardized to 0 mean and 0.5 standard deviations) were entered without interactions. Note that testing interactions between medium, pose, and year is not possible because leg patterns of frogs cannot be photographed out of water without restraint, and we do not have ‘water-restrained’ images from the first year because we came up with this method only for the second year based on our findings from the first year. To avoid the problem of separation in binomial models, we used a Bayesian implementation of linear mixed-effects models with package ‘blme’ in R 4.3.2 (R Core Team, 2023). For defining priors, we followed Gelman’s method as recommended in in Abrahantes and Aerts (2012). We ran this model for FRR_1_, FRR_10_ and FRR_20_, and as a sensitivity analysis also for the corresponding FRRs calculated with ‘corrected ranks’.

To assess observer performance, we calculated FRR_obs_, using the number of matching pairs present among the 10 (in 2023) or 20 (in 2024) highest ranked images on the observer’s computer as denominator in the FRR formula, because these were the images actually searched by the observers. Additionally, we calculated the false acceptance rate (FAR_obs_) for the query results by each observer, which is the number of false matches accepted by an observer divided by the total number of screened image pairs (i.e. number of potential matches screened for each query multiplied by the number of queries assessed), expressed as a percentage.

### Ethical note

This study conforms to Directive 2010/63/EU and was approved by the Ethics Committee of the Plant Protection Institute. All procedures were permitted by the Environment Protection and Nature Conservation Department of the Pest County Bureau of the Hungarian Government (PE/EA/295-7/2018, PE/EA/00270-6/2023, PE-06/KTF/07949-6/2023, PE-06/KTF/00754-8/2022, PE-06/KTF/00754-9/2022). When handling the frogs, we paid attention not to cause harm and unnecessary stress to the animals by keeping handling time as short as possible. We avoided introducing infectious diseases to the studied breeding population by disinfecting all equipment with 96% ethanol before the start of animal handling.

## Results

### Identification success with different photography methods

Regardless of sex, all true matches received rank=1 by HotSpotter in the ‘water-restrained’ databases and the ‘water-restrained subset’ (FRR_1_=0%), while only fewer true matches received rank=1 in all other databases (FRRs ranging from 4 to 50 %; Table 1). Overall, FRR_10_ and FRR_20_ were relatively low for all databases (Table 1). Statistical analyses showed that the usage of both hand-restrained pose and water medium significantly improved FRR with all three rank thresholds, while sex and database size had no significant effect on FRRs (Table 2). For FRR_1_, the methodological changes applied in 2024 resulted in significant improvement compared to 2023, with similar tendencies for FRR_10_ and FRR_20_ (Table 2). All these results were supported by the sensitivity analyses, where ranks were corrected for potential ‘false hotspots’ in the 2024 ‘dry-restrained’ images, with two exceptions: 1) the effect of year was significant for FRR_20_ as well, and 2) both FRR_1_ and FRR_20_ increased with database size, with a similar tendency for FRR_10_ (Table 2).

**Table 2.**
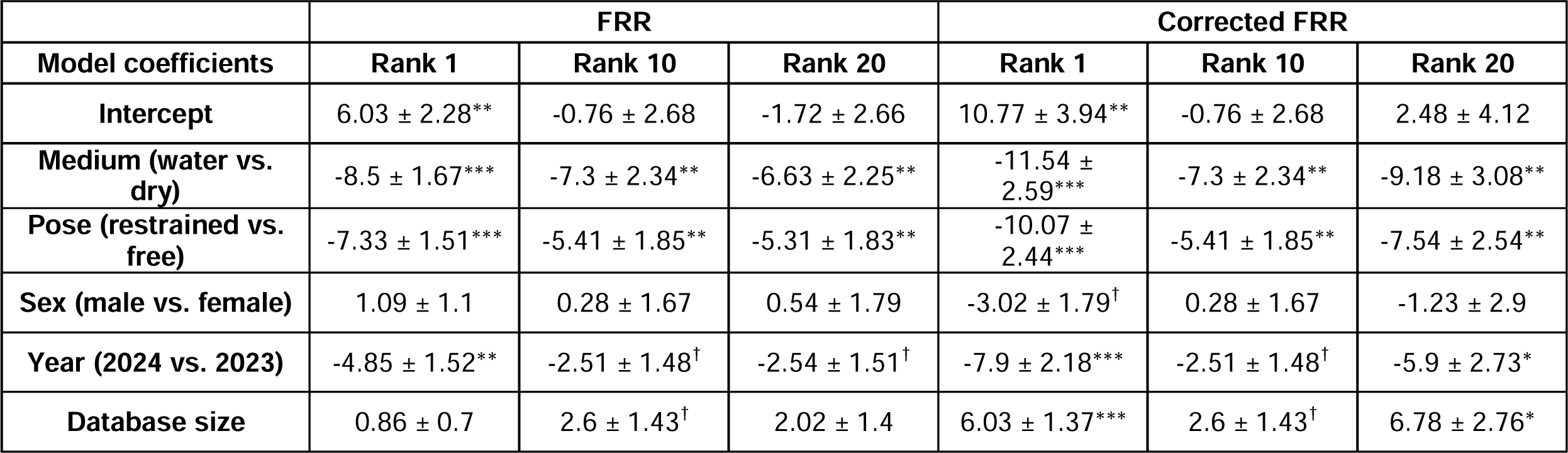
Results of the Bayesian generalized linear mixed-effects models with FRR values calculated either based on the original ranks f or all databases (FRR), or based on corrected ranks in the ‘dry-restrained 2024’ but original ranks in all the other databases (Corrected FRR). Significant effects are denoted by asterisks (*P < 0.05, **P < 0.01, ***P < 0.001) and marginally non-significant effects (0.05 < P < 0.095) are denoted by dagger symbols (†).

|  | **FRR** |  |  | **Corrected FRR** |  |  |
| --- | --- | --- | --- | --- | --- | --- |
| **Model coefficients** | **Rank 1** | **Rank 10** | **Rank 20** | **Rank 1** | **Rank 10** | **Rank 20** |
| **Intercept** | 6.03 ± 2.28** | -0.76 ± 2.68 | -1.72 ± 2.66 | 10.77 ± 3.94** | -0.76 ± 2.68 | 2.48 ± 4.12 |
| **Medium (water vs. dry)** | -8.5 ± 1.67*** | -7.3 ± 2.34** | -6.63 ± 2.25** | -11.54 ± 2.59*** | -7.3 ± 2.34** | -9.18 ± 3.08** |
| **Pose (restrained vs. free)** | -7.33 ± 1.51*** | -5.41 ± 1.85** | -5.31 ± 1.83** | -10.07 ± 2.44*** | -5.41 ± 1.85** | -7.54 ± 2.54** |
| **Sex (male vs. female)** | 1.09 ± 1.1 | 0.28 ± 1.67 | 0.54 ± 1.79 | -3.02 ± 1.79<sup>†</sup> | 0.28 ± 1.67 | -1.23 ± 2.9 |
| **Year (2024 vs. 2023)** | -4.85 ± 1.52** | -2.51 ± 1.48<sup>†</sup> | -2.54 ± 1.51<sup>†</sup> | -7.9 ± 2.18*** | -2.51 ± 1.48<sup>†</sup> | -5.9 ± 2.73* |
| **Database size** | 0.86 ± 0.7 | 2.6 ± 1.43<sup>†</sup> | 2.02 ± 1.4 | 6.03 ± 1.37*** | 2.6 ± 1.43<sup>†</sup> | 6.78 ± 2.76* |

In the ‘water-free 2024’ databases, three out of the four observers successfully identified all matching image pairs when those were provided among the 20 highest-ranked matches by HotSpotter (FRR_obs2-4_=0%), while one observer missed one such match in the ‘water-free 2024’ database of females (FRR_obs1_=2.222%). None of the observers made such mistakes with the ‘water-restrained’ databases, nor with the ‘water-free 2023’ and ‘dry-restrained 2023’ databases (FRR_obs1-4_=0%). Regardless of database type, observers never recorded non-matching images as matching in 2024 (FAR_obs1-4_=0%); despite that two out of the four observers made such mistakes with the ‘water-free 2023’ database type a year earlier (FAR_obs1_=1.176% and FAR_obs2_=0.588%, respectively).

## Discussion

Overall, matching images received significantly better ranks in HotSpotter when frogs were photographed in water instead of out in the dry. We assume that the main reason for this trend was that, when frogs were photographed dry, shadows generated by back muscles and uneven skin surfaces obscured some melanin patterns and induced false ‘hotspots’. (see Supplement Section V; Fig. S3 and Fig. S4). Due to differential light refractions, these shadows vanished when frogs were submerged in water, facilitating melanin-pattern comparison for HotSpotter. In this respect, our findings correspond with a previous report where HotSpotter performed somewhat better with in-water photos than with out-of-water photos of sea turtles (Dunbar et al., 2021). These results show that the choice of medium can matter for the success of photographic ID, and this may be fruitful to consider for a wider range of taxa. More generally, environmental light conditions such as water turbidity for aquatic animals (Gygax et al., 2018; Kelley et al., 2012), or forest versus open-field light for terrestrial animals (Gomez & Théry, 2004), may be worth taking into account when trying to find the best setup for ID-photographing. On the other hand, our results did not support the idea that allowing the animals to take a natural body posture in water would facilitate ID success. Instead, the more-or-less uniform pose of hand-restrained frogs permitted better performance for both HotSpotter and the human observers, once we had eliminated the technical problems associated with dry photography we detected in the first year. Thus, photographing the frogs under water in a standardized hand-restrained position enabled very fast ID whereby the observers had only to check the first-ranked match in HotSpotter to find out the ID of the queried image, and they did so with zero FRR and zero FAR. Nevertheless, after technical improvements from the first to the second year, we found reasonably good ID success with the ‘dry-restrained’ and the ‘water-free’ methods as well, although with these image types we had to check the 10 best-ranked matches of HotSpotter for each query to keep FRR below 10%.

Survival rate and abundance are two key metrics in population monitoring, and to assess how reliable a database is for calculating these, it is essential to calculate FRR and FAR as well. False rejection leads to underestimation of survival rate and overestimation of abundance, while false acceptance has the opposite effects. FRR values calculated in our ‘water-free 2024’ and ‘dry-restrained 2024’ databases were comparable to those reported in the most successful of previous anuran ID studies (Table S1). In contrast, when restrained individuals were photographed in water, all true matches received the highest rank in HotSpotter (FRR_1_=0). To our knowledge, no previous study with similarly large anuran databases demonstrated such high success for photo-based ID. Previously Matthé et al. (2017) found that success rate increases (i.e. FRR decreases) in smaller subset databases when using various other software, and we found qualitatively similar correlation between database size and FRR using HotSpotter. The largest databases used in our study consisted of 426 images, corresponding with the relatively larger database sizes for which FRR or interchangeable metrics were reported in previous anuran studies (Table S1). Furthermore, FRR decreases as the number of images per individual increases in a database (Matthé et al., 2017). Because we included maximum one matching image pair for each individual in our databases, including multiple images would be expected to further improve performance of our methods, especially for the ‘water-free’ databases where different images of the same frog feature different leg postures. In fact, anuran databases with the lowest FRRs published so far contained more than two images of certain individuals (Davis et al., 2020; Matthé et al., 2017; Morrison et al., 2016; Patel & Das, 2020); this contrast further highlights that the zero FRR found here when restrained frogs were photographed in water is outstanding.

The image types we tested here by HotSpotter are expected to be suitable for further platforms as well (Bohnett et al., 2023; Crall, 2020). Observers in general were able to identify matching images captured by either photography method, although FRR_obs_ and FAR_obs_ results taken together from 2023 and 2024 suggest that at least some observers may make mistakes with the ‘water-free’ image type. Previous studies rarely assessed FAR in anuran photo-based identification, but in those few cases where such attempts were made, either no false acceptance was reported (Caorsi et al., 2012; Morrison et al., 2016), or it was lower than 1% (Kim et al., 2017). FAR was similar in our study as well (0% for ‘restrained’ and 0-1% for ‘water-free’ photos).

While in this study we used photographs featuring agile frogs, we expect that the conclusions of our study would stand for various other anuran species as well, due to similarities in melanin-pattern distribution across the body, and behaviour in water (Fig. 1). We propose that photographing anurans in water would be beneficial practice for various monitoring programs, not only because improved melanin-pattern visibility can lead to more reliable ID, but also because these procedures may be less stressful for the frogs than photographing them out of water. Amphibians are most comfortable when their skin is wet, whereas they can remain underwater for much longer than needed for taking a photograph, because they can exchange gas through their skin (Tattersall, 2007). We noticed no specific reaction by agile frogs to being submerged for the ‘water-restrained’ photography (e.g. males continued without interruption the pulsing vocalization that they often use when grabbed by other males or humans). It is likely that letting them move freely in water is even less stressful for them. Therefore, usage of the ‘water-free’ image type may be a preferred choice when working with sensitive species or small juveniles (i.e. to minimise handling time, and to prevent potential injuries when handling small and fragile animals), even at the expense of a somewhat slower or slightly less accurate ID compared to the ‘water-restrained’ method.

In practice, infrastructural limitations and differences in manpower requirement may constrain what photography method can be used in a specific project. For example, while a single person can photograph freely-moving animals alone, photos featuring hand-restrained frogs require the assistance of a second person as well. The usage of images captured by varying cameras and under varying conditions has been identified as a challenge for anuran ID (Aevarsson et al., 2022; Dawson et al., 2021; Kim et al., 2017; Morrison et al., 2016). Here we used a single camera and standard artificial light conditions indoors, which may not be feasible for all studies, but still there is a lot researchers can do to improve image quality. Based on our experiences from the present study, photo-based ID can be improved by avoiding glare caused by direct light exposure, keeping the water clean and choosing appropriate container size and water level (as described in Supplement Section II). Keeping the above viewpoints in mind, applying these photography methods to other species should be fairly straightforward.

All in all, comparison of the ranks received by true matches across different HotSpotter databases demonstrated that 1) photographing frogs in water instead of with towel-dried skin improved ID, and 2) images of hand-restrained frogs in a standardized position submerged in water outperformed the other image types, but 3) photographing agile frogs moving freely in water also demonstrated to be a reasonable option for ID. We propose that the methods developed here, which enable the assessment of patterns across the entire body including legs, can facilitate a refined and more reliable photo-based ID by improving the quality of data compared to conventionally used photography methods. Finally, our results highlight that taking into account the natural conditions under which colour patterns evolved may facilitate photo-based individual identification for animal monitoring.

## Acknowledgements

Part of the funding came from the National Research, Development and Innovation Office of Hungary (K-135016) to V.B. This research was funded in part by the Austrian Science Fund (FWF) [grant ESP 239-B]. For open access purposes, the author has applied a CC BY public copyright license to any author-accepted manuscript version arising from this submission. M.Z. was supported by the EKÖP-24 University Excellence Scholarship Program of the Ministry for Culture and Innovation from the source of the National Research, Development and Innovation Fund, and by the National Research, Development and Innovation Office of Hungary (PD-134241). N.U. was supported by ‘SEH Grant in Herpetology’ and the EKÖP-MATE/2024/25/K university research Scholarship Program of the Ministry for Culture and Innovation from the source of the National Research, Development and Innovation Fund.

We are grateful to Mihály Rusz for his help in both field work and individual identification in HotSpotter, and to Márk Szederkényi, Emese Balogh, Judit Baumann, Gábor Berkei, István Göcző, Dávid Herczeg, Attila Hettyey, Szabolcs Hócza, Beatrix Laczi, András Rotter, Zoltán Simanovszky, Ábris Tóth, János Ujszegi for their participation in the field work.

## Author contributions

This study was conceived, databases were developed and first draft of the manuscript was written by E.N. Statistics were carried out by V.B. and E.N. Field work and photography was carried out by E.N., M.Z., V.B., N.U. and A.K. Individual identification required for creating test databases were carried out by E.N. and N.L. Test databases were manually operated for assessing observer success by N.U., A.K., M.Z. and V.B. All authors reviewed the manuscript and contributed to its final version.

## References

Abrahantes, J. C., & Aerts, M. (2012). A solution to separation for clustered binary data. Statistical Modelling, 12(1), 3–27. 10.1177/1471082X1001200102

Aevarsson, U., Graves, A., Carter, K. C., Doherty-Bone, T. M., Kane, D., Servini, F., Tapley, B., & Michaels, C. J. (2022). Individual identification of the lake Oku clawed frog (*Xenopus longipes*) using a photographic identification technique. Herpetological Conservation and Biology, 17(1), 67–75.

Bendik, N. F., Morrison, T. A., Gluesenkamp, A. G., Sanders, M. S., & O’Donnell, L. J. (2013). Computer-assisted photo identification outperforms Visible Implant Elastomers in an endangered salamander, Eurycea tonkawae. PLoS ONE, 8(3), e59424. 10.1371/journal.pone.0059424

Bohnett, E., Holmberg, J., Faryabi, S. P., An, L., Ahmad, B., Rashid, W., & Ostrowski, S. (2023). Comparison of two individual identification algorithms for snow leopards *(Panthera uncia)* after automated detection. Ecological Informatics, 77(February). 10.1016/j.ecoinf.2023.102214

Bolger, D. T., Morrison, T. A., Vance, B., Lee, D., & Farid, H. (2012). A computer-assisted system for photographic mark–recapture analysis. Methods in Ecology and Evolution, 3(5), 813–822. 10.1111/j.2041-210X.2012.00212.x

Burgstaller, S., Gollmann, G., & Landler, L. (2021). The green toad example: A comparison of pattern recognition software. North-Western Journal of Zoology, 17(1), 96–99.

Canessa, S., Ottonello, D., Rosa, G., Salvidio, S., Grasselli, E., & Oneto, F. (2019). Adaptive management of species recovery programs: A real-world application for an endangered amphibian. Biological Conservation, 236(May), 202–210. 10.1016/j.biocon.2019.05.031

Caorsi, V. Z., Santos, R. R., & Grant, T. (2012). Clip or Snap? An evaluation of toe-clipping and photo-identification methods for identifying individual southern red-bellied toads, *Melanophryniscus cambaraensis*. *South American Journal of Herpetolog*,y7(2), 79–84. 10.2994/057.007.0210

Crall, J. P. (2020). IBEIS: Image based ecological information system. Available at https://github.com/Erotemic/ibeis.

Crall, J. P., Stewart, C. V., Berger-Wolf, T. Y., Rubenstein, D. I., & Sundaresan, S. R. (2013). HotSpotter-Patterned species instance recognition. *Proceedings of IEEE Workshop on Applications of Computer Vision*, January, 230–237. 10.1109/WACV.2013.6475023

Davis, H.-P., VanCompernolle, M., & Dickens, J. (2020). Effectiveness and reliability of photographic identification methods for identifying individuals of a cryptically patterned toad. Herpetological Conservation and Biology, 15(1), 204–211.

Dawson, J., Panter, C. T., & Zeisset, I. (2021). Comparisons of image-matching software when identifying pool frog (Pelophylax lessonae) individuals from a reintroduced population. Herpetological Journal, 31(1), 55–59. 10.33256/31.1.5559

Donnelly, M. A., Guyer, C., Juterbock, J. E., & Alford, R. A. (1994). Techniques for marking amphibians. Measuring and Monitoring Biological Diversity. Standard Methods for Amphibians (Biological Diversity Handbook), 277–284.

Dunbar, S. G., Anger, E. C., Parham, J. R., Kingen, C., Wright, M. K., Hayes, C. T., Safi, S., Holmberg, J., Salinas, L., & Baumbach, D. S. (2021). HotSpotter: using a computer-driven photo-id application to identify sea turtles. Journal of Experimental Marine Biology and Ecology, 535, 151490. 10.1016/j.jembe.2020.151490

Gomez, D., & Théry, M. (2004). Influence of ambient light on the evolution of colour signals: Comparative analysis of a Neotropical rainforest bird community. Ecology Letters, 7(4), 279–284. 10.1111/j.1461-0248.2004.00584.x

Gygax, M., Rentsch, A. K., Rudman, S. M., & Rennison, D. J. (2018). Differential predation alters pigmentation in threespine stickleback (Gasterosteus aculeatus). Journal of Evolutionary Biology, 31(10), 1589–1598. 10.1111/jeb.13354

Jain, A. K. (2007). Are biometric techniques the future of personal identification? Nature, 449, 38–40. 10.1038/449038a

Kelley, J. L., Phillips, B., Cummins, G. H., & Shand, J. (2012). Changes in the visual environment affect colour signal brightness and shoaling behaviour in a freshwater fish. Animal Behaviour, 83(3), 783–791. 10.1016/j.anbehav.2011.12.028

Kim, M. Y., Borzée, A., Kim, J. Y., & Jang, Y. (2017). Treefrog lateral line as a mean of individual identification through visual and software assisted methodologies. Journal of Ecology and Environment, 41(1), 42. 10.1186/s41610-017-0060-1

Lama, F. Del, Rocha, M. D., Andrade, M. Â., & Nascimento, L. B. (2011). The use of photography to identify individual tree frogs by their natural marks. South American Journal of Herpetology, 6(3), 198–204. 10.2994/057.006.0305

Luedtke, J. A., Chanson, J., Neam, K., Hobin, L., Maciel, A. O., Catenazzi, A., Borzée, A., Hamidy, A., Aowphol, A., Jean, A., Sosa-Bartuano, Á., Fong G, A., de Silva, A., Fouquet, A., Angulo, A., Kidov, A. A., Muñoz Saravia, A., Diesmos, A. C., Tominaga, A., … Stuart, S. N. (2023). Ongoing declines for the world’s amphibians in the face of emerging threats. Nature. 10.1038/s41586-023-06578-4

Lukacs, P. M., & Burnham, K. P. (2005). Review of capture-recapture methods applicable to noninvasive genetic sampling. Molecular Ecology, 14(13), 3909–3919. 10.1111/j.1365-294X.2005.02717.x

Matthé, M., Sannolo, M., Winiarski, K., Spitzen - van der Sluijs, A., Goedbloed, D., Steinfartz, S., & Stachow, U. (2017). Comparison of photo-matching algorithms commonly used for photographic capture–recapture studies. Ecology and Evolution, 7(15), 5861–5872. 10.1002/ece3.3140

Morrison, T. A., Keinath, D., Estes-Zumpf, W., Crall, J. P., & Stewart, C. V. (2016). Individual identification of the endangered Wyoming Toad *Anaxyrus baxteri* and implications for monitoring species recovery. Journal of Herpetology, 50(1), 44–49. 10.1670/14-155

Patel, N. G., & Das, A. (2020). Shot the spots: a reliable field method for individual identification of *Amolops formosus*(Anura, Ranidae). Herpetozoa, 33, 7–15. 10.3897/HERPETOZOA.33.E47279

Price, A. C., Weadick, C. J., Shim, J., & Rodd, F. H. (2008). Pigments, patterns, and fish behavior. Zebrafish, 5(4), 297–307. 10.1089/zeb.2008.0551

R Core Team. (2023). R: A language and environment for statistical computing. R ver. 4.3.2. R Foundation for Statistical Computing, Vienna, Austria. http://www.r-project.org.

Rojas, B. (2017). Behavioural, ecological, and evolutionary aspects of diversity in frog colour patterns. Biological Reviews, 92(2), 1059–1080. 10.1111/brv.12269

Rojas, B., Lawrence, J. P., & Márquez, R. (2023). Amphibian Coloration: Proximate Mechanisms, Function, and Evolution. In Evolutionary Ecology of Amphibians (Issue April 2023, pp. 219–258). CRC Press. 10.1201/9781003093312-12

Sarasola-Puente, V., Gosá, A., Oromí, N., Madeira, M. J., & Lizana, M. (2011). Growth, size and age at maturity of the agile frog *(Rana dalmatina)* in an Iberian Peninsula population. Zoology, 114(3), 150–154. 10.1016/j.zool.2010.11.009

Tattersall, G. J. (2007). Skin breathing in amphibians. In W. C. Aird (Ed.), Endothelial Biomedicine (Issue 10, pp. 85–91). Cambridge University Press. 10.1017/CBO9780511546198.010

The RawTherapee Team. (2022). RawTherapee 5.9. Accessed from https://rawtherapee.com/. https://rawtherapee.com/

