## Supplement for "Put the frog in water: simple methods for improving individual identification as demonstrated with the agile frog"

#### I. Individual identification

In 2023, individuals were identified by manually comparing images of frogs leaving the pond with images of same-sex frogs arriving to the pond. Images of the arriving individuals were grouped by SVL to reduce the effort required for manual identification (each leaving animal was compared with all arriving animals with similar SVL (allowing for ± 3 mm inconsistency). The results of manual identification were used for creating HotSpotter databases of individuals with known identity, including images of leaving frogs that either did or did not have matching pairs among the images captured at arrival. All males were identified manually, and completeness of the recapture results was confirmed by running queries for each capture event in HotSpotter later. A subset of female images was also manually compared within SVL categories (the thus-identified capture events were used for creating HotSpotter databases in 2023, see the next section), but completed HotSpotter databases were used later for identifying all capture events in females as well, and these databases were continued the next year.

In 2024, we created four 'full databases' (two for each sex) in HotSpotter that contained images (one or more) from each capture event across 2023 and 2024. One database type consisted of images predominantly featuring hand-restrained agile frogs (photographed either dry or in water), while another database contained images mainly featuring agile frogs moving freely in water. Queries were performed in HotSpotter for each capture event in 2024 in all databases by one of three people in total (two authors and a volunteer; but free-moving databases were assessed by a single person, the database creator), who manually assessed the 20 highest-ranked matches per query and assigned an animal identifier (the 'name' variable in HotSpotter) to each image that they deemed matching. Using an R code developed under R version 4.2.3 (R Core Team, 2023), we extracted data from the csv files saved by HotSpotter and created comprehensive tables for further processing and manual comparison in Excel (Microsoft Office 365). Using these tables, we manually double-checked images of all those capture events where results differed between the two databases (i.e. matching image from another capture event was indicated in one, but not in the other database). After these double checks, we deemed individuals to be successfully identified.

#### II. Practical guidelines for anuran photography

We fine-tuned photographing methods over the course of two agile frog breeding seasons, and during this time we noted some mistakes and tricks that should be kept in mind when photographing frogs in water. First, it's important to keep the water clean, because debris (e.g. pieces of soil and shed skin) and bubbles accumulating in the water can obscure melanin patterns on the images. Second, it is important to choose a container where even the largest individuals can comfortably stretch their legs (e.g. in agile frogs, females can be substantially larger than males). Third, choosing an optimal water level is important: when photographing hand-restrained frogs, the water level should be high enough to enable complete submerging of the body even for the largest individuals such as gravid females. For photographing freely-moving frogs, taller container is required, and the water level should not be close to the top of the container's wall, otherwise the frogs may jump out. Although we did not specifically collect data on frog behaviour, we noted that when the water level was low (i.e. 10-20 centimetres from the top of the container's wall, but higher than the length of the forelimbs of the animals), agile frogs were easier to photograph, because they either stayed underwater or floated at water level, usually automatically taking postures suitable for photographing. Even when the above aspects are considered, one should be patient when photographing free-moving frogs, should capture multiple images, and may need to gently initiate posture changes by hand if needed, to ensure suitability for ID. We also noticed that photography skills may make greater difference in image suitability when animals are not restrained.

For photographing animals out of water, it is important to minimize shadows caused by uneven skin surfaces and glare caused by wetness that may remain on the gently dried skin. Here we used standard artificial light conditions indoors. However, we acknowledge that photographing frogs in the field will limit researchers' options in this respect. In any case, we advise against using the camera's flashlight. Using a photo box with battery-powered light source mounted in a standard way to avoid reflections may be a more suitable solution (Kim et al., 2017).

#### III. Detailed methods of image selection

In 2023, the male databases created for analyses consisted of 98 individuals and a total of 108 images (one per capture event) each: 88 at entering the pond, and 20 at leaving it, representing all male capture events during the study period. However, manual identification revealed that one individual was captured three times: leaving, entering and leaving the pond again, respectively; therefore, we subsequently decided to leave out all query results for this individual from the analyses, to ensure that each queried image had only one possible match in each database. We captured more than twice as many females as males in 2023, but due to the significant effort required for manual identification (often based on multiple images per capture event), and also because in-hand photos were available for only a limited number of females, we used a smaller subset of known-identity females for creating the 'dry-restrained 2023' and 'water-free 2023' datasets of females: images from 80 capture events were included (63 at entering the pond and 17 at leaving it; out of which 11 leaving images had a true matching image in the database), representing a total of 69 individuals.

To create the 'water-restrained 2024' and 'water-free 2024' databases, we removed images from the 'full databases' by the criteria detailed below. First, only images captured in 2024 were included (this was necessary due to methodological changes in the photography techniques between 2023 and 2024). Second, we only kept images that were captured on a day (or partial day) when greater photography errors were not typical. This means that photographs were mostly taken by the same person within each day, and methodological requirements were either not clear to certain photographer-handler teams or they lacked necessary photography skills (e.g. they captured images in steep angles, did not avoid reflections stemming from direct light exposure, or photographed animals in dirty water); the images taken by these teams were omitted. Third, only images of those capture events were kept in the test databases that were photographed by both the 'water-restrained' and 'water-free' methods, to ensure database comparability. Following these guidelines, we excluded 7 full days and 2 partial days from the total of 31 days when agile frogs were captured in 2024. Lastly, from images remaining after the above three filtering steps, we kept only one image per capture event (the image deemed most suitable for identification overall), and for animals that were captured multiple times, only the first two capture events of those still remaining were kept in the test databases. As a result, the test databases for both 'water-restrained' and 'water-free' images were identical for each sex regarding both the represented capture events (and hence individuals) and the number of images: 426 images in the male and 199 images in the female databases, featuring 374 and 153 individuals, respectively. As a last step, we removed the animal identifiers (the 'name' variable in HotSpotter) for all 'chips'.

We created smaller 'dry-restrained 2024' databases, because the 'dry-restrained' photo type was only available for fewer capture events. Furthermore, we created an identically sized 'water-restrained subset' database for each sex, to compare these two image types at the same database size. We did this because database size may affect FRR, and the main databases were approximately twice as large for males as for females, while the opposite was true for the 'dry-restrained 2024' databases. The 'dry-restrained 2024' and 'water-restrained subset' databases consisted of 186 images for females (representing all capture events in the main databases, except for 13 when 'dry-restrained' images were not captured), and 101 images for males (representing all capture events with true matches present in the main databases, except for 3 when ‘dry-restrained’ images were not captured).

#### IV. Note on sharing HotSpotter databases

All databases were created by the researcher performing manual identification (database creator), and each database was tested independently by two out of four other researchers (observers). In 2023, two researchers worked with the male databases and two other researchers worked with the female databases. In contrast in 2024, each observer assessed one image type for one sex, and another image type for the other sex. All four observers were blind to the identities of the photographed frogs, and they had no previous experience with HotSpotter before 2023. In 2023 we noted that queries ran in the same HotSpotter database can result in somewhat different scores and different ranks on different computers (apparently only if queries have not yet been performed in the shared database version), and we consciously accounted for this in 2024. Databases saved before query running on the database creator’s computer were shared with the observers via pendrive without file compression (but in 2023, some observers received the databases via Google Drive after zip compression). Subsequently, in each database, queries were run in the same order on the database creator’s computer as well as on the computers of two observers to enable comparison of scores and ranks across three computers.

#### V. Distribution of ‘hotspots’

Using the ‘sample’ function in R (R Core Team, 2023), we randomly chose 10 male and 10 female individuals from those which were photographed with all three methods in 2024. We manually assessed the distribution of matched ‘hotspots’ across the body on image pairs captured with the ‘dry’, in-water ‘hand-restrained’ and ‘free-moving’ methods. We categorized these ‘hotspots’, and assumed that those denoting melanin patterns were most reliable for identification (see categories on Fig. S3). ‘Hotspots’ on patches that could be described as shade differences rather than clear melanin patterns fell into a separate, less reliable category (see Fig. S4 for illustration of longer-term changes of such traits).

We found that on most images in the ’dry-restrained 2024’ database, some ‘hotspots’ were assigned to shadows rather than, or in addition to melanin patterns. Such misleading ‘hotspots’ were not recognized elsewhere, because shadows were not visible on images photographed in water (except on one ‘water-free’ image where the eyes emerged above water level, but it did not generate a ‘hotspot’). The handler’s fingertips, which were not cropped from the ‘chips’ in the ’water-restrained’ and ’dry-restrained’ databases, were rarely included in ovals denoting ‘hotspots’, and in most of those cases their involvement seemed rather coincidental (e.g. majority of the ovals also contained clear melanin patterns). The number of ‘hotspots’ on melanin patterns, as well as overall, was usually notably lower on the ‘water-free’ images. This may be attributed to differences in body postures and photo angles between the query and matching ‘chips’, and to the fact that the size of melanin dots was relatively smaller within the area denoted by the ‘chip’ on the ‘water-free’ images compared to the relative size of the same melanin dots on the ‘chip’ of either the ‘water-restrained’ or ‘dry-restrained’ images of the same individual. With other words, ‘chips’ in the ‘water-free’ databases were wider and less zoomed in due to the free leg postures (see panels A and B on Fig. 3 for reference).

### VI. Supplementary tables and figures

**Table S1. False rejection rates (FRR) in previous computer-assisted individual identification studies with anuran amphibians.** Published results by three thresholds are shown: matching was considered successful if at least one matching image 1) received the highest rank ($FRR_1$), 2) was among the 10 highest-ranked images ($FRR_{10}$) or 3) was among the 20 highest-ranked images ($FRR_{20}$)

| Study | Species | N images | N image/ individual | Software | $FRR_1$ (%) | $FRR_{10}$ (%) | $FRR_{20}$ (%) |
|---|---|---|---|---|---|---|---|
| Matthé et al (2017) [1] | *Bombina variegata* | varying: 500-4000 | varying (2-10) | multiple | 2.3-19.6 | 1.2-11.4 | - |
| | | 4063 | varying (2-10) | multiple | 3.1-19.6 | 1.7-11.4 | - |
| Patel & Das (2020) | *Amolops formosus* | 301 | varying (2-4) | HotSpotter | 5.6 | - | - |
| Caorsi et al (2012) [1] | *Melanophryniscus cambaraensis* | 492 | - | Wild-ID | 26 | - | - |
| Davis et al (2020) [1] | *Rhinella diptycha* | 109 | varying (2-3 or more) | Wild-ID | - | - | 0? [3] |
| Dawson et al (2021) [1] | *Pelophylax lessonae* | 465 | - | multiple | 45.9-59.2 | - | - |
| Burgstaller et al (2021) [2] | *Bufotes viridis* | 200 | 2 | multiple | c.a. 40-60 | - | - |
| | | 200 | 2 | HotSpotter | c.a. <10 | - | - |
| Morrison et al (2016) | *Anaxyrus baxteri* | 130 | - | HotSpotter and Wild-ID | - | - | 56.6-76.2 |
| Aevarsson et al (2022) | *Xenopus longipes* | 48 | 2 | Wild-ID | - | - | 0-17 |
| | | 20 | 2 | | - | - | 100 |
| Kim et al (2017) | *Dryophytes japonicus* | - | - | Wild-ID | - | - | 25 |
| | | 213 ? | - | | - | - | 100 |

[1] FRR was derived from the reported 'success rate' (100-'success rate').

[2] FRR was derived from 'correctly identified images %' shown on Figure 3 (100-'correctly identified images %').

[3] Cut-off criteria for 'success' are not clear, so the '100% success rate' might reflect other than true matches present between ranks 1 and 20.

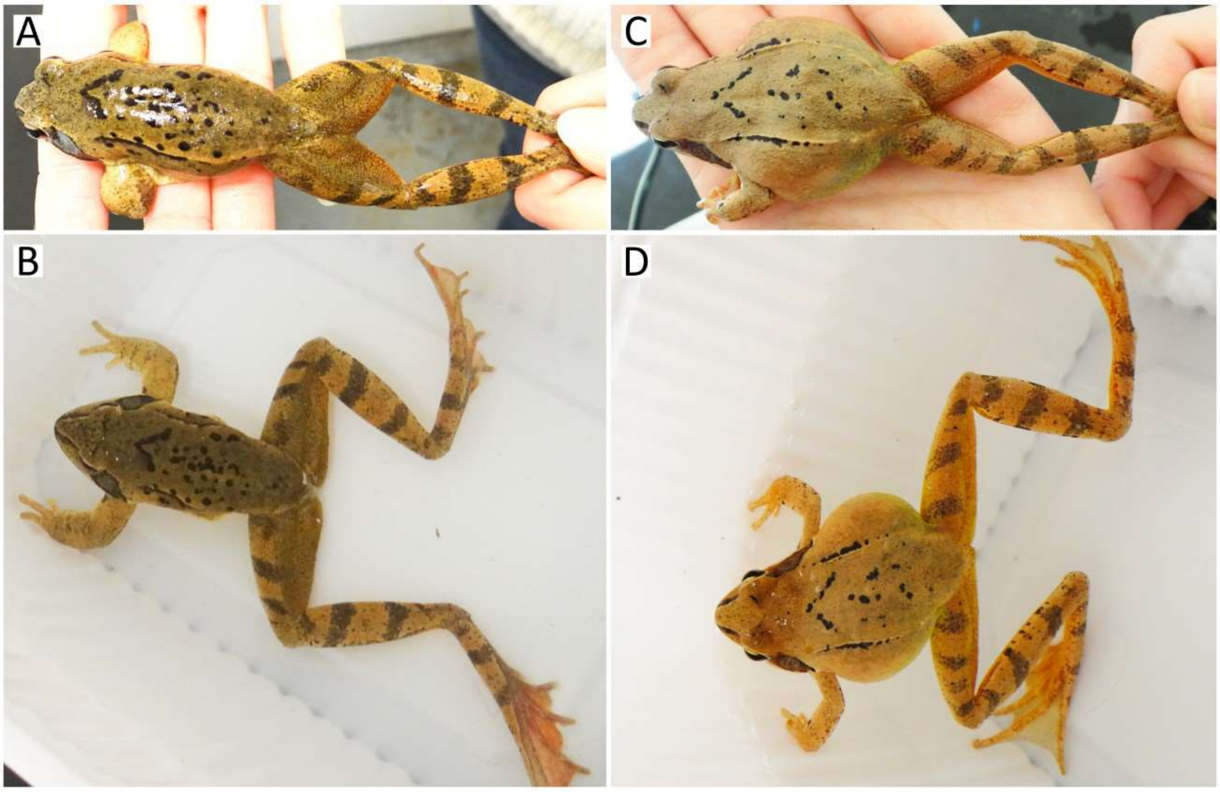

**Fig. S1. Examples of a male (A, B) and a female (C, D) individual photographed in hand on dry (A, C) and freely in water (B, D) in 2023.** Direct exposure to artificial ceiling light caused strong glare on some individuals despite being dried by paper towel (A), and some frogs took postures that made photographing more difficult, such as standing vertically in the water (D) instead of floating or laying horizontally (B). Note the small bubbles and pieces of dirt in the water over the frog body (B, D).

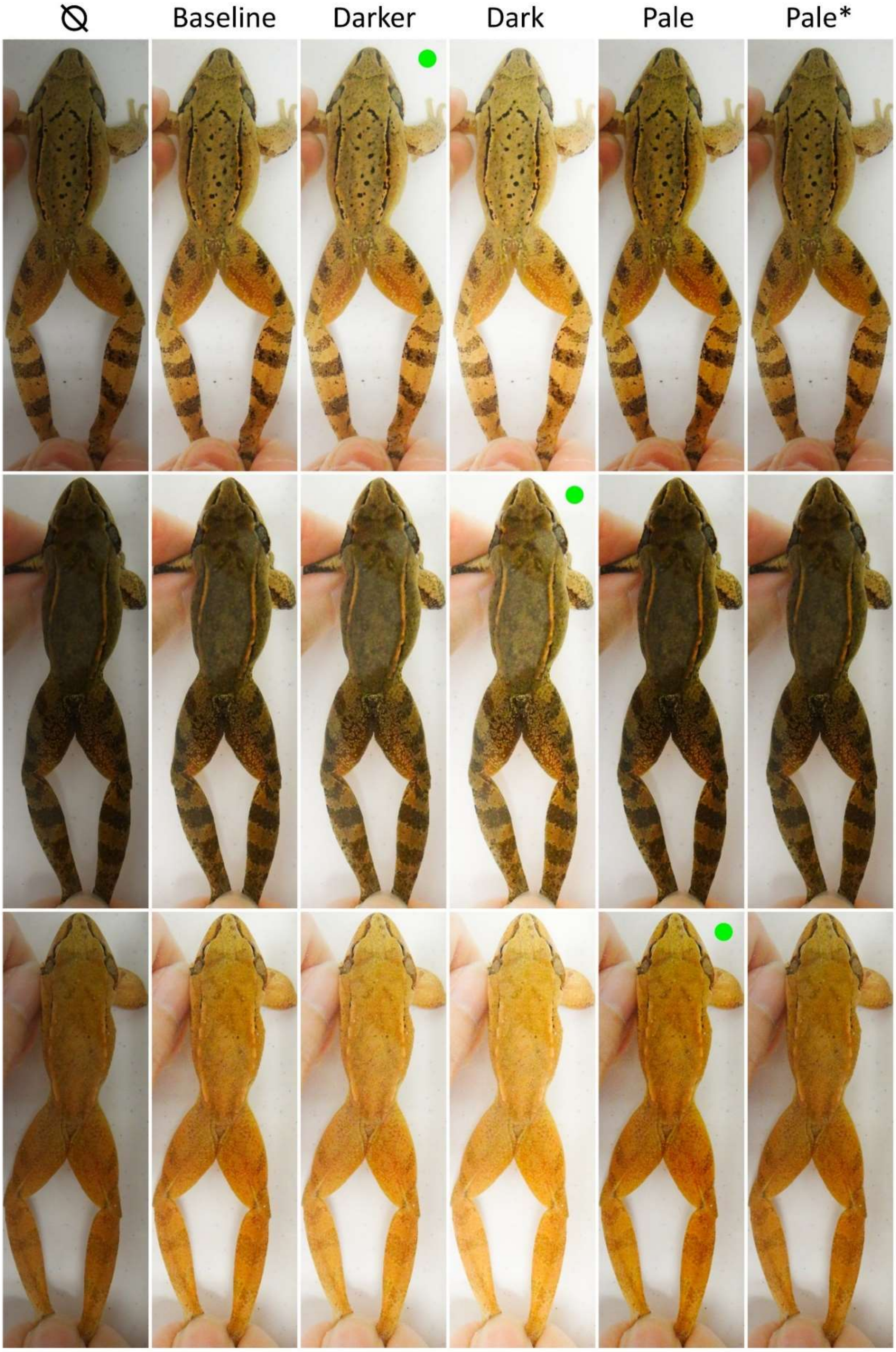

**Fig. S2. Examples of different skin tone and melanin-pattern strength combinations and the results of applying different exposure curves in RawTherapee in 2024.** The setting deemed best in terms of melanin-pattern visibility for each photo is marked with a green circle. Short name for each curve is shown on top (the leftmost version is the original photo); the exposure curve marked with asterisk was created for the special case when a pale animal is featured on an unusually bright photo. Also, note the lack of bubbles and pieces of dirt in the water over the frog body.

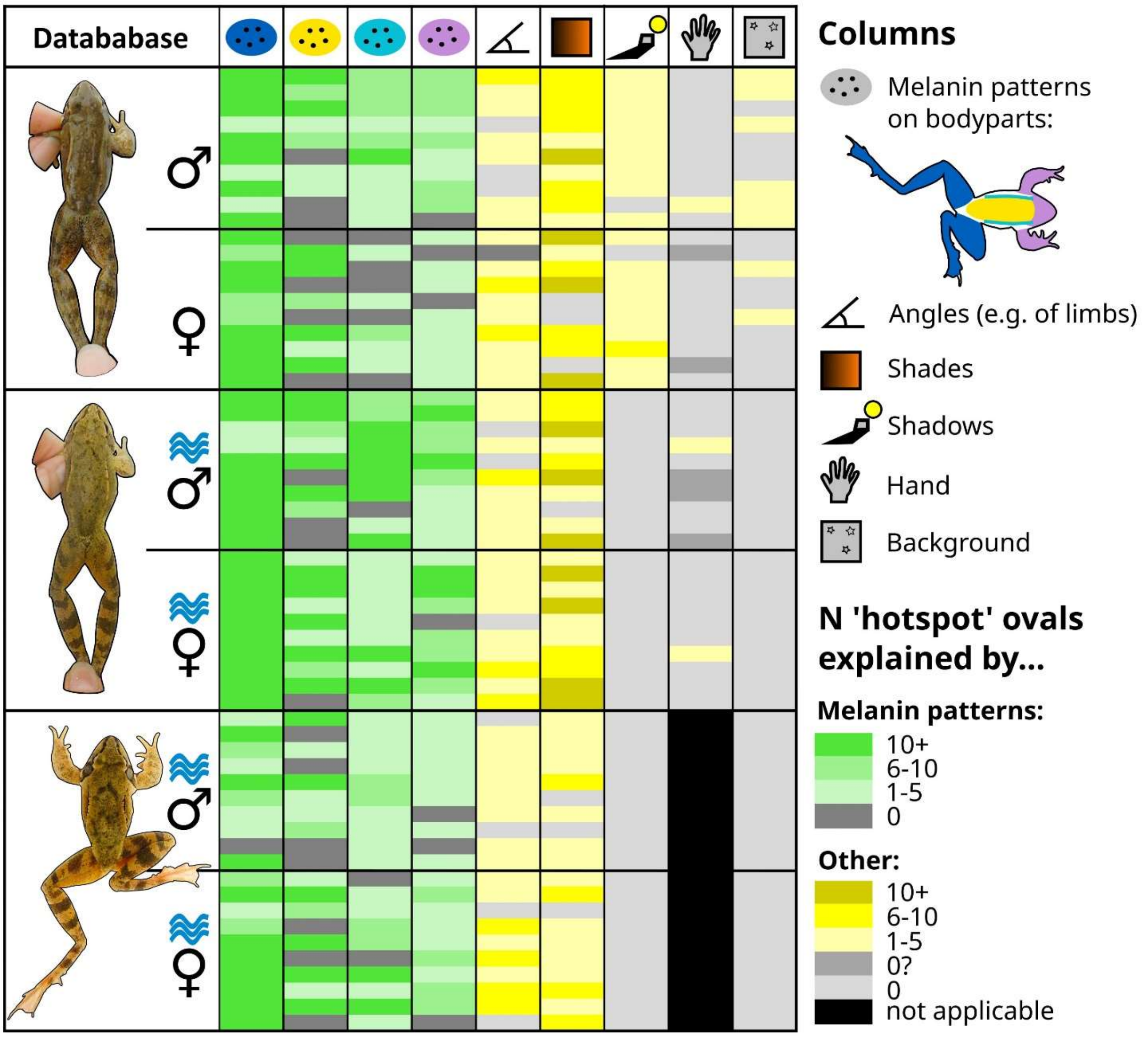

**Fig. S3. Visual assessment of 'hotspot' distribution in different databases.** The order of ten randomly-chosen male and female individuals (one row per individual) is the same across database types 'dry-restrained' (top), 'water-restrained' (center) and 'water-free' (bottom), respectively. Ovals displayed by HotSpotter denoting 'hotspots' were categorized by a single human observer.

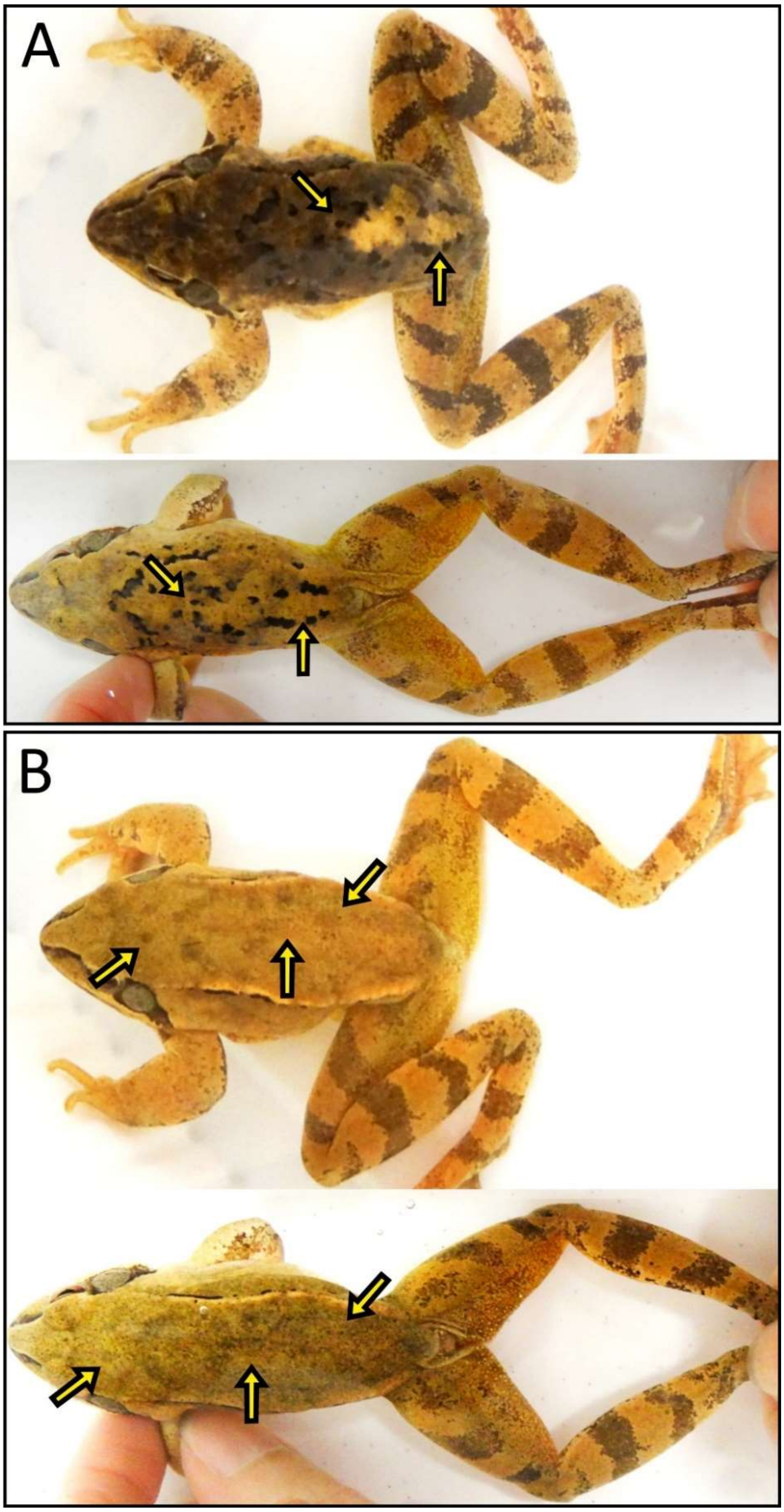

**Fig. S4. Illustration of strong (A) and mild (B) changes of distinct-shade spots on the back.** Within each panel, the upper and lower image features the same animal in 2023 and 2024, respectively.

#### References

Aevarsson, U., Graves, A., Carter, K. C., Doherty-Bone, T. M., Kane, D., Servini, F., Tapley, B., & Michaels, C. J. (2022). Individual identification of the lake Oku clawed frog (*Xenopus longipes*) using a photographic identification technique. *Herpetological Conservation and Biology*, *17*(1), 67–75.

Burgstaller, S., Gollmann, G., & Landler, L. (2021). The green toad example: A comparison of pattern recognition software. *North-Western Journal of Zoology*, *17*(1), 96–99.

Caorsi, V. Z., Santos, R. R., & Grant, T. (2012). Clip or Snap? An evaluation of toe-clipping and photo-identification methods for identifying individual southern red-bellied toads, *Melanophryniscus cambaraensis*. *South American Journal of Herpetology*, *7*(2), 79–84. https://doi.org/10.2994/057.007.0210

Davis, H.-P., VanCompernolle, M., & Dickens, J. (2020). Effectiveness and reliability of photographic identification methods for identifying individuals of a cryptically patterned toad. *Herpetological Conservation and Biology*, *15*(1), 204–211.

Dawson, J., Panter, C. T., & Zeisset, I. (2021). Comparisons of image-matching software when identifying pool frog (Pelophylax lessonae) individuals from a reintroduced population. *Herpetological Journal*, *31*(1), 55–59. https://doi.org/10.33256/31.1.5559

Kim, M. Y., Borzée, A., Kim, J. Y., & Jang, Y. (2017). Treefrog lateral line as a mean of individual identification through visual and software assisted methodologies. *Journal of Ecology and Environment*, *41*(1), 42. https://doi.org/10.1186/s41610-017-0060-1

Matthé, M., Sannolo, M., Winiarski, K., Spitzen - van der Sluijs, A., Goedbloed, D., Steinfartz, S., & Stachow, U. (2017). Comparison of photo-matching algorithms commonly used for photographic capture–recapture studies. *Ecology and Evolution*, *7*(15), 5861–5872. https://doi.org/10.1002/ece3.3140

Morrison, T. A., Keinath, D., Estes-Zumpf, W., Crall, J. P., & Stewart, C. V. (2016). Individual identification of the endangered Wyoming Toad *Anaxyrus baxteri* and implications for monitoring species recovery. *Journal of Herpetology*, *50*(1), 44–49. https://doi.org/10.1670/14-155

Patel, N. G., & Das, A. (2020). Shot the spots: a reliable field method for individual identification of *Amolops formosus* (Anura, Ranidae). *Herpetozoa*, *33*, 7–15. https://doi.org/10.3897/HERPETOZOA.33.E47279

R Core Team. (2023). *R: A language and environment for statistical computing. R ver. 4.3.2.* R Foundation for Statistical Computing, Vienna, Austria. http://www.r-project.org.
